# Modular assembly of mitochondrial complex III from a dynamic matrix domain pool

**DOI:** 10.64898/2026.11.05.764608

**Authors:** Jorge Jimenez-Niebla, Roshan Jha, Christophe Wirth, Jon V. Busto, Antoine Feignier, Miguel Cosenza-Contreras, Katharina Ruoff, Manuel Dorer, Riccardo Babic, Friedel Drepper, Claudine Kraft, Pitter F. Huesgen, Nils Wiedemann, Wei-Chun Kao, Carola Hunte

**Affiliations:** Institute of Biochemistry and Molecular Biology, ZBMZ, Faculty of Medicine, University of Freiburg, Freiburg, 79104, Germany; Faculty of Biology, University of Freiburg, Freiburg, 79104, Germany; Spemann Graduate School of Biology and Medicine (SGBM), University of Freiburg, Freiburg, 79104, Germany; Biochemistry and Functional Proteomics, Faculty of Biology, University of Freiburg, Freiburg, Germany; CIBSS - Centre for Integrative Biological Signalling Studies, University of Freiburg, Freiburg, 79104, Germany

## Abstract

Mitochondrial complex III is a central component of the cellular energy metabolism. It is an obligate dimer of catalytic and accessory subunits that are encoded by two genomes. How its large matrix domain, formed by the core proteins Qcr1 and Qcr2, is built and joined to the membrane part and during assembly is not known. Here we show that Qcr1 and Qcr2 preassemble independently of complex III into matrix domain modules. Cryo-EM structures from a cardiolipin-deficient yeast strain and of recombinant protein resolved single, double and triple matrix domains, which are linked by the C-terminal tail of Qcr1. In the mature complex III, the same tail binds cytochrome *b* and cardiolipin. The modules form a dynamic pool, in which assembly species interconvert and exchange subunits. Mutations at the module interfaces, including mimics of human disease variants, reshape this pool and deletion of the Qcr1 tail impairs respiratory growth. Substitution of a tyrosine residue linked to parkinsonism exposes a conserved cysteine residue to oxidation. A cryo-EM structure of an assembly intermediate shows a cap of two matrix domains bound to a monomeric membrane module of cytochrome *b*, Qcr7, Qcr8 and cytochrome *c*_1_. We propose that complex III receives its matrix domain from a dynamic pool of prebuilt modules, which organizes the dimerization of the complex.

## Introduction

Mitochondrial respiratory complex III (cytochrome *bc*_1_ complex) transfers electrons from ubiquinol to cytochrome *c* and couples this reaction to proton translocation across the inner mitochondrial membrane. It is an obligate homodimer at the center of the respiratory chain (*1-3*). In yeast, each protomer contains three catalytic subunits, cytochrome *b* (cyt *b*, Cob) with two b-type heme groups, cyt *c*_1_ with a c-type heme bound, and the Rieske protein Rip1 with a 2Fe-2S cluster, as well as seven accessory subunits (*1*). Cyt *b* is mitochondrially encoded, whereas the other subunits are of nuclear origin. The two largest accessory subunits, the core proteins Qcr1 and Qcr2, form most of the matrix domain. Assembling such a complex is demanding. Its subunits are encoded by two genomes, redox cofactors must be inserted in a defined order, and partly assembled complexes can be harmful (*4, 5*). Respiratory complex I meets this challenge by assembling from preformed modules and the use of several assembly factors (*6*).

Complex III, in contrast, is thought to assemble in a step-wise manner on the membrane. For the early assembly steps, binding of the Cbp3-Cbp6 chaperone to nascent cyt *b* was described (*7, 8*). The studies suggest as next steps hemylation of cyt *b* and binding of Qcr7 and Qcr8. In the late assembly phase, the AAA-ATPase Bcs1 inserts the folded Rip1 (*9*). The steps in between are poorly defined. Biochemical studies described the presence of core proteins in a dimeric intermediate with cyt *b*, cyt *c*_1_ and Qcr6 (*10, 11*). Recent cryo-EM structures show a membrane bound assembly intermediate that is already a dimer (*12*).There are no assembly factors known for these intermediate steps. How the large matrix domain is assembled, delivered and coupled to dimerization is therefore unknown.

Transient assembly intermediates are difficult to capture. We reasoned that the lack of cardiolipin which is known to be important for complex III function and binds at defined sites of the complex (*13, 14*), might slow down assembly and enrich such states. In the cardiolipin-deficient yeast strain *crd1*Δ (*15*), we found free heterooligomers of Qcr1 and Qcr2 that are not assembled with complex III. Here we show these are prebuilt matrix domain modules, which form a dynamic pool in mitochondria and *in vitro*. Interface mutations, including mimics of human disease variants, reshape this pool, and one of them exposes a conserved cysteine residue to oxidation. Finally, a cap of two matrix domains binds a monomeric early assembly intermediate, before complex III dimerizes.

## Results

### Qcr1 and Qcr2 form free matrix domain modules not associated with complex III

We purified a soluble Qcr1/Qcr2 assembly from *crd1*Δ mitochondria and determined its structure by cryo-EM at 2.5-Å resolution (Fig. 1A, Table S1). It is a heterododecamer of six Qcr1 and six Qcr2 subunits. Each Qcr1 pairs tightly with one Qcr2 to form a core module (CM; mean buried area 1511 Å^2^), and two CMs pair through their Qcr2 subunits (mean buried area 940 Å^2^). This CM pair has the arrangement of the matrix domain of mature complex III (Fig. 1 C and D). The dodecamer is therefore a trimer of matrix domains, which we call CM_Hexa_.

**Fig. 1.**
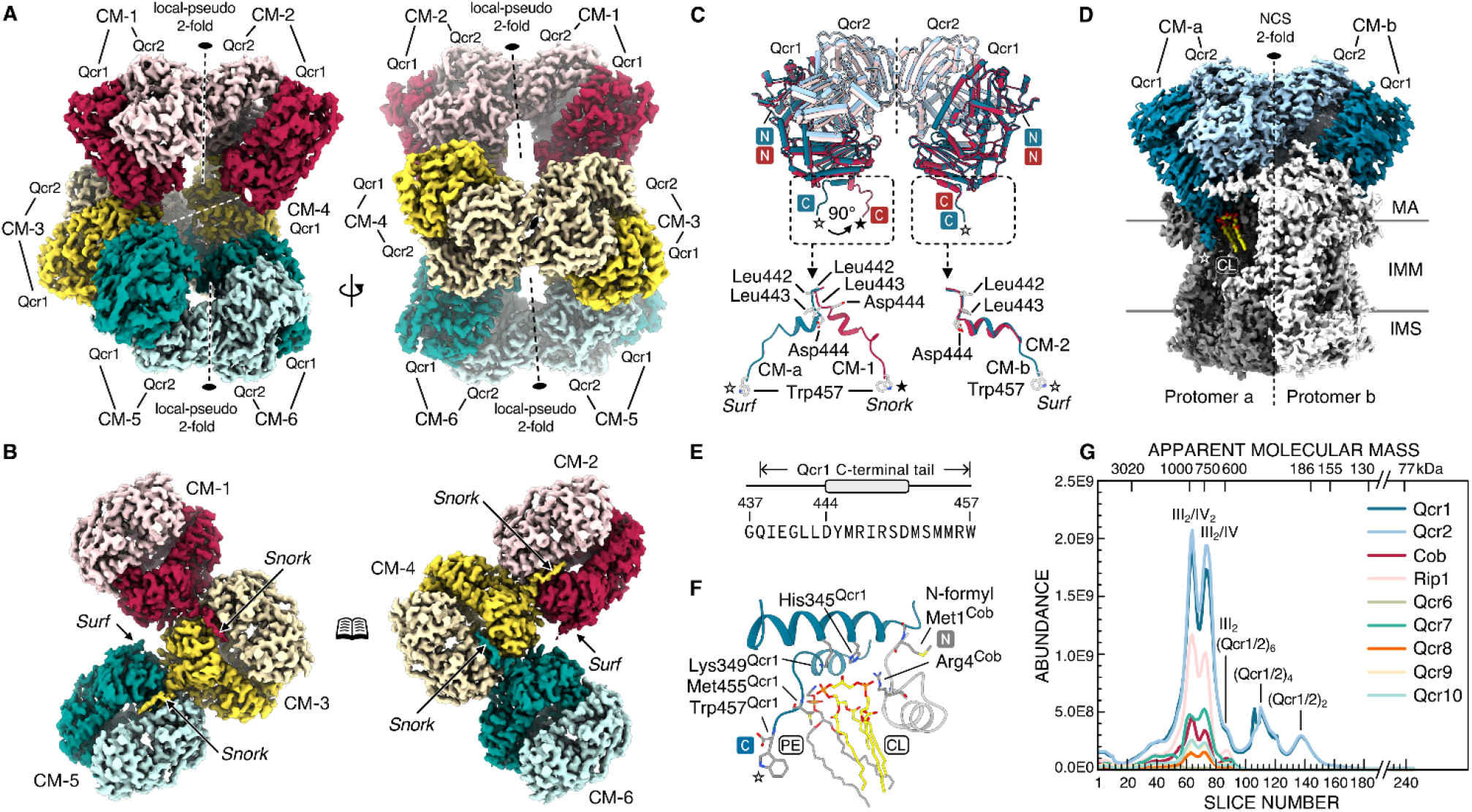
Trimer of preassembled matrix domains of complex III. **(A)** Front (left) and back (right) view of cryo-EM structure of Qcr1/Qcr2 heterododecamer from *crd1*Δ, organized in six core modules (CM) of Qcr1/Qcr2 (CM_Hexa_, Table S1) (**B**) Snorkeling of the Qcr1 C-terminal tail (*Snork*) into adjacent CM connects the domains in an intertwined manner (open-book view), leaving two termini surface exposed (*Surf*) in assembly-competent orientation. (**C)** Complex III matrix domain highlighted in blue as in X-ray structure of complex III ((D), Table S1) and single domain of CM_Hexa_ (red) are structurally very similar including the Qcr1 tail in *Surf* position, whereas it tilts by 90° in *Snork* position (**E**) Scheme and amino acid sequence of Qcr1 tail. (**F**) In the mature complex III (D), the tail of Qcr1 binds the N-terminal formyl-methionine of the central catalytic subunit cyt *b* (Cob), together forming a binding site for cardiolipin bound at the surface of the quinone exchange cavity next to a phosphatidyl-ethanolamine (PE). (**G**) Distinct hetero-oligomers of Qcr1/Qcr2 are present in the mitochondrial high-resolution complexome MitCOM from wild-type (WT) yeast (*16*). MA, matrix; IMM, inner mitochondrial membrane; IMS, intermembrane space.

The three matrix domains are associated in a distinct manner. Notable is the position of the C-terminal tail of Qcr1 (residues Q438-W457) (Fig. 1, B and E). In the central domain, both Qcr1 tails reach into the Qcr1/Qcr2 cleft of a neighboring domain, a position we call *Snork*. Each outer domain inserts only one tail, and its second tail is positioned on the surface (*Surf*). CM_Hexa_ thus carries two exposed tails that are free for further contacts (Fig. 1B).

Each matrix domain of CM_Hexa_ closely matches the matrix domain in the X-ray structure of complex III (Cα RMSD 0.45 Å, averaged from superimposed Qcr1) including the Qcr1 tail in *Surf* position (Fig. 1, C and D). In *Snork* position, the C-terminal tail is turned by about 90° from its helix onwards (Fig. 1C). In the mature complex III, the tail has a defined role. There, it binds the N-terminal formyl-methionine of cyt *b* (Cob), and together they form the binding site for a cardiolipin molecule at the quinone exchange cavity, next to a phosphatidylethanolamine (Fig. 1F). The same peptide thus links the matrix domains to each other in the free assembly, but links the matrix domain to cyt *b* and to a cardiolipin in complex III. Loss of cardiolipin may destabilize this contact site in mature complex III and thus favor the free assembly, which would explain why CM_Hexa_ was abundant in *crd1*Δ.

Matrix domain modules are not restricted to the mutant. The high-resolution complexome of wild-type yeast mitochondria (MitCOM) (*16*) contained distinct Qcr1/Qcr2 assembly species besides complex III and its supercomplexes, with apparent masses that fit assemblies of matrix domains (Fig. 1G).

### The modules form a dynamic pool in mitochondria and *in vitro*

Qcr1 and Qcr2 were abundant in the soluble fraction of both wild-type and *crd1*Δ mitochondria (Fig. 2A), for which native PAGE resolved them as two distinct heterooligomers (Fig. 2B). In the membrane fraction, they were part of complex III (III_2_) and of the small (III_2_/IV) and large (III_2_/IV_2_) supercomplexes (Fig. 2C). Complex III still assembles without cardiolipin, but free complex III was more abundant than the supercomplexes in *crd1*Δ, the reverse of wild type (Fig. 2C), in line with the known role of cardiolipin for supercomplex stability (*14*). The Qcr1/Qcr2 heterooligomers co-purified with complex III in ion exchange chromatography, but could be also eluted separately with a salt gradient (Fig. 2D). Complexome analysis of *crd1*Δ resolved all complex III subunits in the membrane fraction and two large soluble Qcr1/Qcr2 heterooligomers (Fig. 2E).

**Fig. 2.**
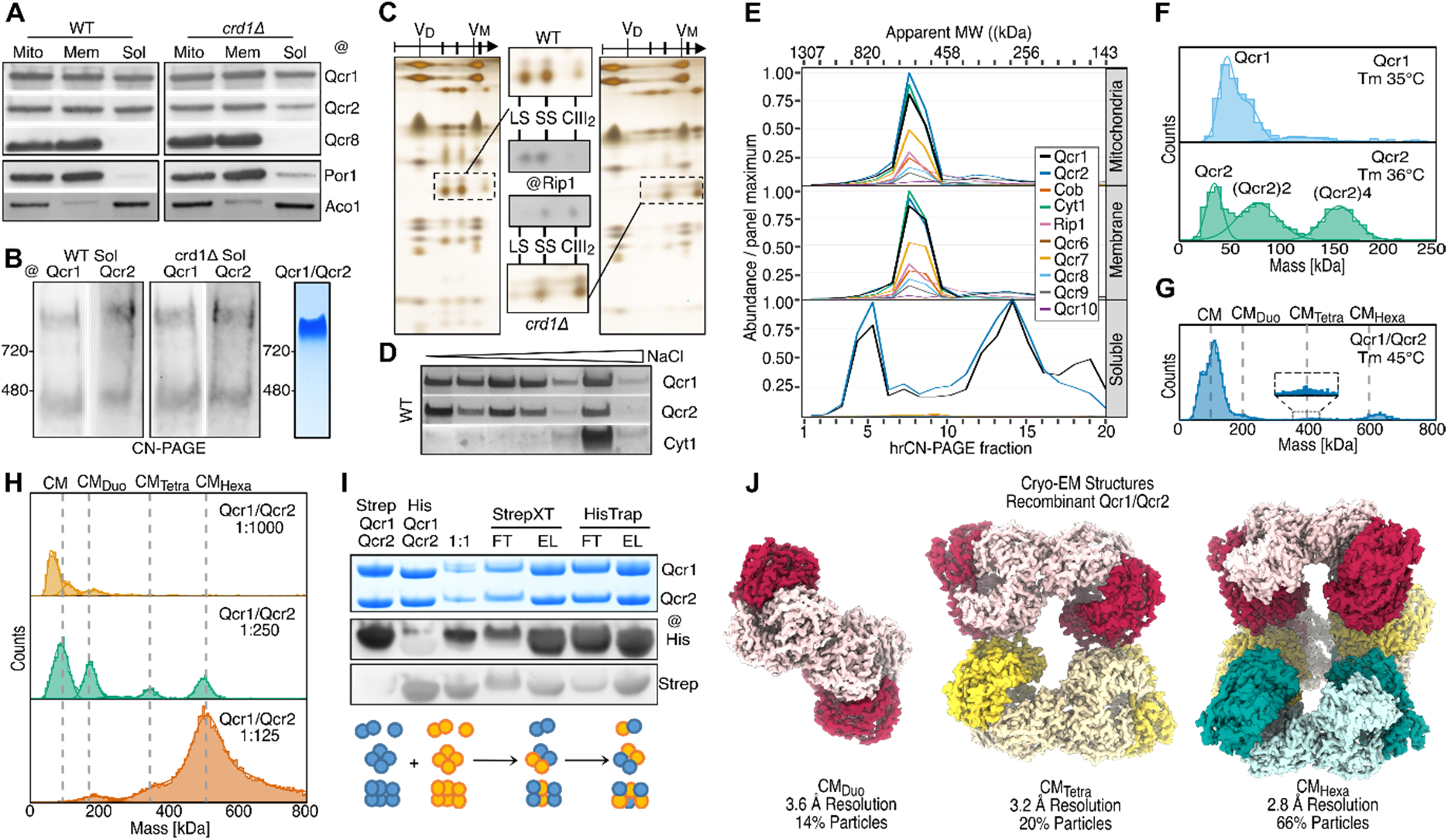
Qcr1/Qcr2 heterooligomers are abundant in mitochondria, self-assemble and form a dynamic pool of preassembled matrix domains. **(A)** Qcr1 and Qcr2 are abundant in the soluble fraction of wild type and *crd*1Δ. (**B**) In these fractions, they are resolved as two distinct heterooligomeric species by native PAGE. Recombinant Qcr1/Qcr2 self-assemble when co-expressed in *E. coli* cultures. (**C**) Qcr1 and Qcr2 are also part of mature complex III (III_2_) and of small (SS) and large (LS) supercomplex (2D-PAGE). Complex III is more abundant in *crd1*Δ than supercomplexes in contrast to the distribution in WT. (**D**) Qcr1 and Qcr2 co-purify with complex III in ion-exchange chromatography but can be separated by a salt gradient. (**E**) Section of complexome analysis of whole mitochondria as well as mitochondrial membrane and soluble fraction of *crd1*Δ separated by CN-PAGE resolved complex III with all subunits in membrane and whole mitochondrial fraction and assigned apparent masses to Qcr1/Qcr2 heterooligmers in the soluble fraction. (**F**) Mass photometry showed strong and little oligomerization of recombinant Qcr2 and Qcr1, respectively. (**G**) Co-expression of Qcr1 and Qcr2 resulted in distinct heterooligomers of CM alone as well single (CM_Duo_), double (CM_Tetra_) and triple (CM_Hexa_) matrix domains, with increased thermal stability compared to subunits alone. Concentrating the sample in native PAGE resulted in one distinct species resembling of higher mass. (**H**) The oligomeric species form a dynamic pool that can be interconverted in concentration dependent manner as shown by microfluidic mass photometry (MassFluidix). (**I**) Qcr1/Qcr2 heterooligomers exchange subunits. (**J**) Cryo-EM analysis of recombinant Qcr1/Qcr2 resolved structures of CM_Duo_, CM_Tetra_ and CM_Hexa_, with the latter being the most abundant component (Table S1).

Qcr1 and Qcr2 do not need other mitochondrial factors to form these modules. Using co-expression in *Escherichia coli* cell cultures, they formed heterooligomers of a size similar to the largest native species in CN-PAGE (Fig. 2B). Alone, Qcr2 oligomerized strongly and Qcr1 only weakly as shown by mass photometry (Fig. 2F). Together, they formed a single CM as well as single (CM_Duo_), double (CM_Tetra_) and triple (CM_Hexa_) matrix domains, with a higher thermal stability than the subunits alone (Fig. 2 F, and G).

The modules are not fixed end products. Their distribution shifted with protein concentration and the species interconverted as shown by microfluidic mass photometry (MassFluidix) indicating a dynamic pool of the distinct assembly species (Fig. 2H). To probe for exchange of subunits, we mixed differently tagged preparations of co-expressed Qcr1/Qcr2, and recovered both tagged proteins together, which shows that the modules exchange subunits (Fig. 2I). Cryo-EM analysis of recombinant Qcr1/Qcr2 resolved structures of CM_Duo_, CM_Tetra_, and CM_Hexa_, with CM_Hexa_ as the most frequent species (Fig. 2J). Qcr1 and Qcr2 thus form a self-assembling, interconverting pool of matrix domain modules, dominated by CM_Hexa_.

### Disease-related interface mutations reshape the pool and expose Cys312 to oxidation

Overall, the contacts within CM_Hexa_ match those of the mature matrix domain (Fig. S2). We identified key residues at the Qcr1/Qcr2 and Qcr2/Qcr2 interfaces of CM_Hexa_, which provide stabilizing contacts and are conserved in sequence and structure between yeast and human subunits (Fig. S1), and are linked to disease (Fig. 3A). At the Qcr2/Qcr2 interface, R152 forms a cation-π interaction with Y364 and a polar interaction with D366 of the neighboring Qcr2. These contacts are central to two patches of cation-π interactions and electrostatic interactions along the interface at the symmetry axis. At the Qcr1/Qcr2 interface, Y288 of Qcr1 stabilizes a loop that contacts Qcr2 and the C-terminal tail of a neighboring Qcr1 in *Snork* position. The human equivalents, R183 in UQCRC2 and Y314 in UQCRC1, are mutated in complex III deficiency and in parkinsonism with polyneuropathy, respectively (*17, 18*). We studied alanine substitutions, the disease mimics R152W and Y288S, and a deletion of the C-terminal Qcr1 tail (ΔCT).

**Fig. 3.**
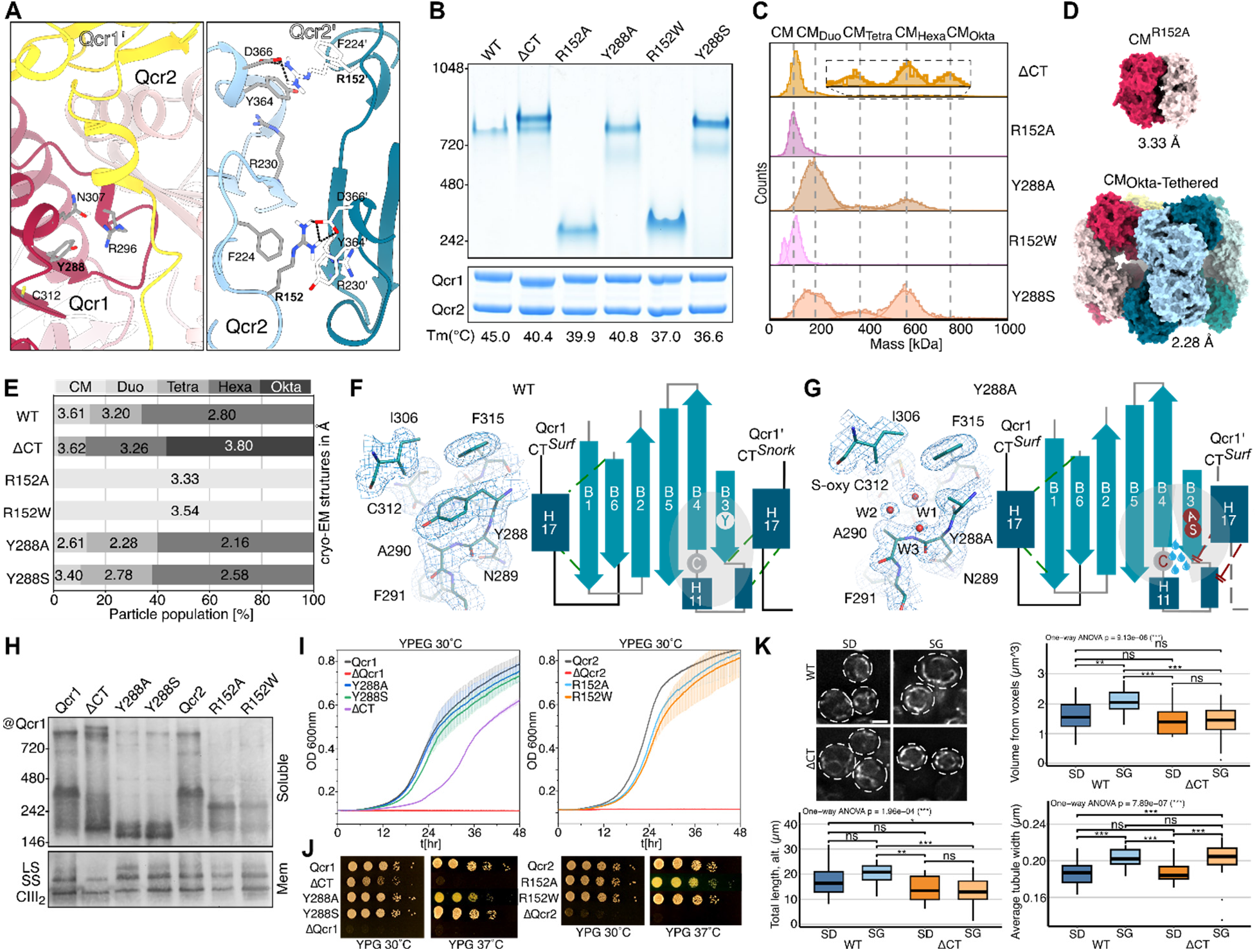
Mutations perturbing the Qcr1/Qcr2 and the Qcr2/Qcr2 interface affect the heterooligomer pool, mitochondrial morphology, and respiratory growth with substitution Y288A causing oxidation of Cys312 of Qcr1. (**A**) Key interface residues are conserved in yeast and man (Fig. S1) including Y288 of Qcr1 at the Qcr1/Qcr2 interface (left panel) and R152 of Qcr2 at the Qcr2/Qcr2 interface. R152 is central to two patches of polar interactions (cation-π, ion-pair, H-bond) at the dimer interface (right panel). (**B, C**) Alanine substitution, mimics of patient mutations (Y288S, R152W) and deletion of Qcr1 tail (ΔCT) in recombinant co-expressed Qcr1/Qcr2 reduce thermal stability compared to WT for all variants, impair hetero-oligomerization for R152A/W probed by CN-PAGE (**B**) and mass photometry (**C**) and cryo-EM analysis (**E**). R152A/W disrupt the dimer interface and enabled cryo-EM structure determination of CM alone (**D**), upper panel (Table S1). ΔCT shifts the heterooligomeric pool to larger assemblies including the CM_Octa_ (C,E) with four matrix domains. (**D**) CM_Octa_ could be enforced by a double cysteine mutation and resolved in a cryo-EM structure (lower panel, Table S1). (**E**) Cryo-EM structures of Qcr1/Qcr2 variants highlighting the relative abundance of the type of Qcr1/Qcr2 heterooligomers (Table S1) (**F**,**G)** Y288A perturbs the Qcr1 ß-sheet, resulting in a water filled cavity at Cys312, its oxygenation, and changing the Qcr1 tail of the adjacent Qcr1 to *Surf* rather than *Snork* position compared to WT, as resolved in cryo-EM structures of recombinant CM_Hexa_ of WT (left panel) and variants (right panel, Y288A,). (**H**) In yeast, ΔCT, Y288A/S, and R152A/W destabilized the Qcr1/Qcr2 heterooligomers. ΔCT substantially impairs respiratory growth (**I, J**) and (**K**) affects mitochondrial morphology.

The mutations affected the pool in two ways. All variants lost thermal stability (Fig. 3B). R152A and R152W broke the Qcr2/Qcr2 contact and impaired module formation, as seen by native PAGE, mass photometry and cryo-EM (Fig. 3B, C, E). Both shifted the pool towards the single CM, which enabled its structure determination (Fig. 3D, E). Loss of the Qcr1 tail had the opposite effect: ΔCT shifted the pool to larger assemblies, including CM_Octa_ with four matrix domains (Fig. 3C, E), which we stabilized with an engineered double cysteine (Qcr1 N171C/Qcr2 R62C) for structure determination (Fig. 3D). The tail therefore prevents the formation of the CM_Octa_ and keeps the Qcr1/Qcr2 pool dynamic, consistent with its role in the *Snork* position.

The Y288 variants revealed a redox switch. In CM_Hexa_ of Y288A, loss of the tyrosine side chain opened a water-filled cavity at the ß-sheet that contains residue 288 (Fig. 3F, G). This cavity extends to Cys312, of which the side chain was oxidized as resolved in the cryo-EM structure of the variant (Fig. 3G). Mass spectrometry confirmed sulfenic and sulfinic acid at Cys312 in recombinant and native protein. Cys 312 is conserved in sequence and structure in human UQCRC1 (Cys338) (Fig. S1). The substitution also moved the neighboring Qcr1 tail from *Snork* to *Surf* position (Fig. 3F). A single interface residue substitution thus links oxidation of a buried cysteine to the way modules are joined.

These mutations also affect cells. We complemented *qcr1*Δ and *qcr2*Δ strains with wild-type or mutant Qcr1 and Qcr2, respectively. R152A, R152W, Y288A and Y288S reduced the higher order heterooligomers in BN-PAGE (Fig. 3H). Moreover, the deletion of the Qcr1 tail impaired respiratory growth (Fig. 3 I, J) and altered the mitochondrial network (Fig. 3K). In yeast, the Qcr1 tail, the key element that shapes the module pool, is therefore needed for normal respiratory function, and the disease-related Y288 variant leads to oxidation of a buried cysteine residue.

### A prebuilt matrix cap docks onto an early assembly state of complex III before dimerization

We next asked where the modules enter the assembly line. In *crd1*Δ, we identified an intermediate that was smaller than complex III (Fig. 4A). Its cryo-EM structure at 2.7-Å resolution contains a single copy each of cyt *b*, Qcr7, Qcr8 and cyt *c*_1_ forming a membrane module (Fig. 4B). On the matrix side, this membrane module carries a CM_Tetra_ cap with two complete matrix domains. Rip1, Qcr6, Qcr9 and Qcr10 are absent.

**Fig. 4.**
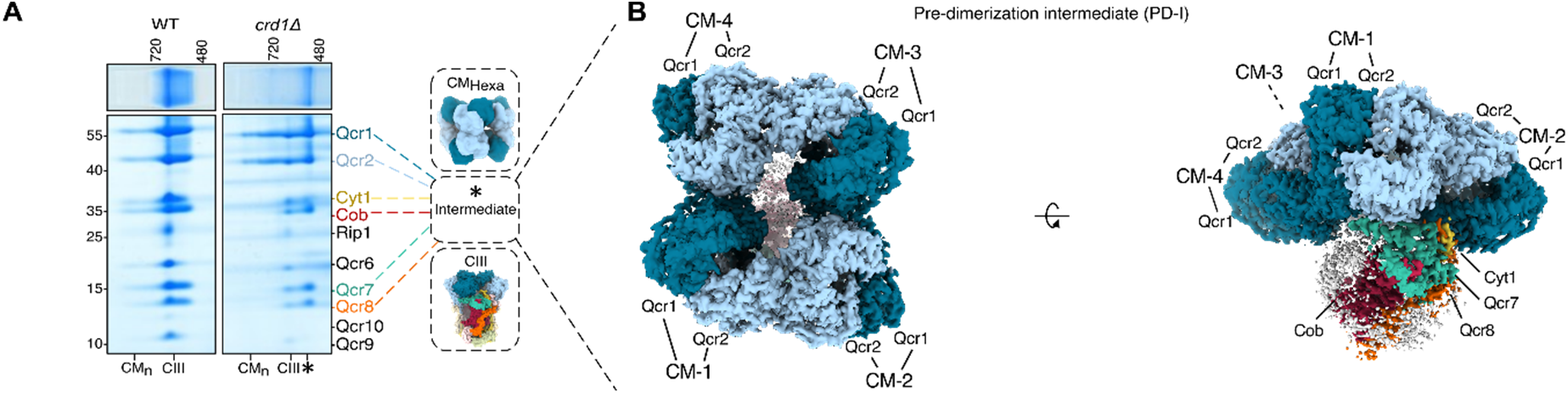
Pre-dimerization intermediate of complex III holds a surplus matrix domain. **(A)** Pre-dimerization intermediate identified in ion-exchange enriched mitochondrial membrane fractions from *crd1*Δ. (**B**) Cryo-EM structure (Table S1) of pre-dimerization intermediate purified from *crd1*Δ revealed a CM_Tetra_ cap docked onto the early assembly state of complex III with cyt *b*, Qcr7, Qcr8 and cyt *c*_1_.

The membrane part of this intermediate is a monomer, whereas the matrix part contains more than a dimer. One CM of the first matrix domain binds the cyt *b*, Qcr7, Qcr8, and cyt *c*_1_ module as in mature complex III (Fig S2). The second CM of this domain has no membrane partner yet, and the second matrix domain is surplus. All complex III assembly intermediates resolved so far have a dimeric membrane part (*12*). Our structure shows that the matrix domain is not added to a dimeric membrane core. Instead, it arrives as a prebuilt dimeric unit, before the membrane part dimerizes and before Qcr6 joins.

## Discussion

Our data support a model in which complex III is built from two modules that form in different compartments (Fig. 5). The membrane module of cyt *b*, Qcr7, Qcr8, and cyt *c*_1_ forms in the inner membrane. The matrix module forms from imported Qcr1 and Qcr2 in the matrix, as a dynamic pool of interconvertible oligomers of preassembled matrix domains. CM_Hexa_ shields-assembly competent surfaces. It can shed a CM_Tetra_ unit, which docks onto the membrane module (Fig. 4). As the cap already carries a complete dimeric matrix domain, its free half offers a docking site for a second membrane module to bind. Release of the surplus domain at this step would complete the dimeric architecture. Thus, the matrix side may organize dimerization. This model fits to the obligate dimeric architecture of complex III (*1, 2*). Our intermediate precedes the dimeric intermediates described recently, in which the intermembrane space domain folds one protomer at a time (*12*).

**Fig. 5.**
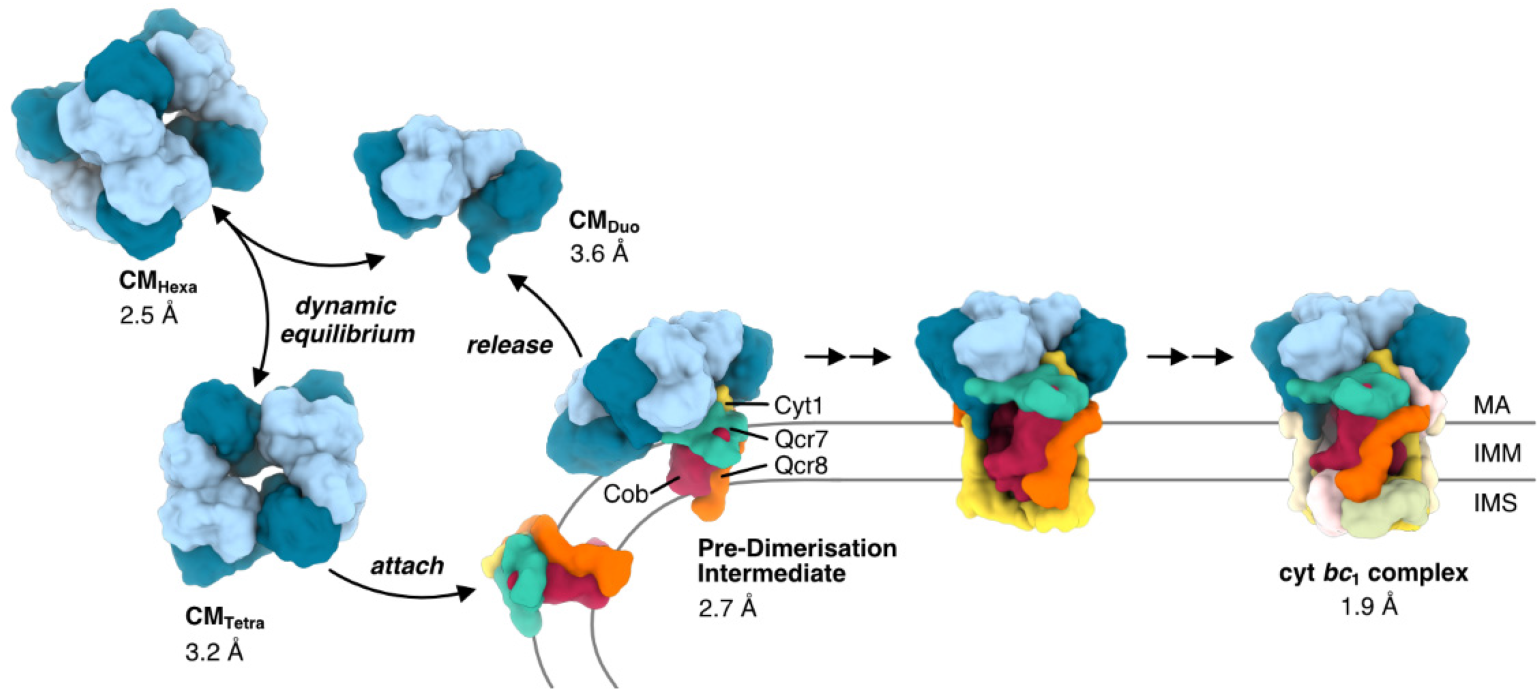
Heterooligomers of Qcr1/Qcr2 form a dynamic pool of preassembled matrix domains that deliver the cap of the pre-dimerization intermediate of complex III. CM_Hexa_ can shed CM_Tetra_ to cap the early assembly state of complex III with cyt b, Qcr7, Qcr8 and cyt *c*_1_. Cryo-EM structures of intermediates determined are indicated in the scheme with resolution given in Å.

The Qcr1 tail may act as a checkpoint along this path. It has three states: *Snork*, which links domains in the pool and prevents the formation of the assembly-impaired CM_Octa_; *Surf*, which is free for assembly; and the anchor state, in which it binds to the cyt *b* and cardiolipin in the mature complex (Fig. 1). The anchor state requires this cardiolipin site. This would explain why the pool and the pre-dimerization intermediate accumulate in *crd1*Δ,. It also suggests that the role of cardiolipin for complex III is not limited to the stability and enzyme activity of the mature complex (*13, 14*), but already starts during assembly, in line with the link between OXPHOS assembly and cardiolipin remodeling (*19*).

A pool of prebuilt modules has clear benefits. It can buffer the supply of imported core proteins against the tightly controlled synthesis of mitochondrially encoded cyt *b* (*5*). It can keep unassembled subunits protected and ready for assembly. It can also deliver a complete matrix domain to the membrane in one step without a dedicated assembly factor, which would explain why no factor has been found for the middle assembly steps.

The pool is also a point open to regulation. Variants that mimic human disease mutations shift its distribution. At the Qcr1/Qcr2 interface, they make Cys312 accessible to reactive oxygen species and change the position of the Qcr1 tail that controls the pool dynamics. Such mutations may therefore act on the assembly of complex III rather than on its catalysis. As incomplete respiratory complexes can be harmful (*4*), oxidation of modules that wait for assembly could feed the redox state of the organelle back into the assembly line.

Complex III has long been seen as the example of a respiratory complex that assembles by sequential addition of subunits. Our results show that it relies on a preformed and dynamic module, similar to the modular assembly of complex I (*6*). Reservoirs of interchangeable subassemblies, of which the distribution is set by interfaces, lipids and redox state, may be a common strategy to build large membrane redox machineries in a robust way.

## Materials and Methods

### Strains and growth conditions

*S. cerevisiae* strains used in this study were grown in YP medium (1% yeast extract and 2% Bacto peptone) supplemented with 3% glycerol (YPG), 2% galactose (YPGal), or 2% glucose (YPD), at 19–23°C with shaking at 130 rpm. Growth plates were prepared by addition of 3% (wt/vol) agar to media. To assess yeast growth strains were cultured overnight in SD-Trp at 30°C. On the next day, cells were diluted to an OD_600_ of 0.3 and cultured for 3h to reach mid-exponential phase. Cells corresponding to 1 mL of OD_600_ = 1.0 were harvested and washed twice in sterile dH_2_O to remove residual medium. Finally, a 10-fold serial dilution was performed, 3 µL of each dilution were plated onto YPD/YPG plates and incubated at 30°C or 37°C.

Growth assays were performed using an EPOCH2 microplate reader (BioTek), and data were collected for up to 48 h. *E. coli* strains DH5α and BL21 were grown in LB medium supplemented with the appropriate antibiotics at 37°C and at 200 rpm, unless otherwise stated. Amino acid substitutions in Qcr1 and Qcr2 were introduced by site-directed mutagenesis using the pETDuet plasmid system, with plasmids carrying the genes encoding for the corresponding mature protein sequences as PCR templates. Primers harboring the desired substitutions were used for PCR amplification. An N-terminal Twin-Strep tag was inserted to facilitate purification. The resulting PCR products were treated with DpnI (NEB) for 1 h at 37°C to remove the template DNA, purified using the QIAquick PCR Purification Kit (Qiagen), and assembled using Gibson Assembly Master Mix (NEB), according to the manufacturer’s instructions. The reaction products were transformed into competent DH5α *E. coli* cells and incubated overnight at 37°C. Single colonies were picked and inoculated into LB medium containing carbenicillin. Plasmids were isolated using the QIAprep Spin Miniprep Kit (Qiagen), and constructs carrying the desired sequence variations were identified by sequencing.

### Purification of complex III, pre-dimerization intermediate and Qcr1/Qcr2 heterooligomers from *S. cerevisiae*

Yeast cells from the BY4741 *crd1*Δ and wild-type strain were grown essentially as described previously (*20*) but in YP-Gal media. Submitochondrial particles were isolated as in previous studies (*1*). Complex III was purified by ion-exchange chromatography using a NaCl elution gradient (self-packed DEAE-Sepharose Fast Flow column (Cytiva)) (*1*), followed by a Heparin purification with NaCl elution (HiTrap Heparin HP column (Cytiva)). Qcr1/Qcr2 heterooligomers and the pre-dimerization intermediate were co-purified.

### Recombinant production and purification of subunits Qcr1 and Qcr2

Plasmids encoding the recombinant proteins were transformed into competent BL21 *E. coli* cells. Transformants were cultured overnight at 37°C with shaking at 200 rpm, diluted into fresh medium to an OD_600_ of 0.1, and incubated at 30°C with shaking at 200 rpm. Protein expression was induced by adding IPTG to a final concentration of 0.5 mM when the cultures reached an OD_600_ of 0.6. Cells were harvested after 24 h of expression. The cell pellets were resuspended in buffer W (50 mM Tris-HCl, pH 8.0, 150 mM NaCl, and 1 mM EDTA) and lysed using a cell disruptor at 25 kpsi. Protein purification was performed on an AKTA pure system (Cytiva). Filtered supernatant was applied to a StreptacinXT 4flow column (iba Lifesciences) pre-equilibrated with buffer W. Protein-containing fractions were analyzed by SDS-PAGE, and relevant fractions were pooled and further purified by size-exclusion chromatography (SEC) using a Superdex 200 HiLoad 26/600 column (Cytiva). Elution fractions were analyzed by SDS-PAGE; fractions with the highest purity were pooled, and protein concentration was determined using a BCA protein assay (Thermo). For analytical SEC, purified proteins were applied to a Superdex 200 Increase 10/300 GL column (Cytiva) equilibrated in buffer W.

### Isolation of mitochondria

Yeast cells were grown in YPG or YPGal to an OD_600_ of 2 or 20, respectively, for mitochondrial isolation. Cells were harvested by centrifugation at 4,000 rpm for 8 min, washed, and resuspended in 1.2 M sorbitol and 20 mM potassium phosphate-HCl (pH 7.4) containing Zymolyase-20T (Amsbio; 3 mg per g wet cell weight). After incubation for 30 min at 24°C, the spheroplasts were homogenized on ice using a glass-Teflon homogenizer (25 strokes) in homogenization buffer (0.6 M sorbitol, 10 mM Tris-HCl, pH 7.4, 1 mM EDTA, and 0.2% fatty acid-free bovine serum albumin (Sigma)). Crude mitochondrial fractions were collected from the homogenate by differential centrifugation (2 min at 1,500 rpm, followed by 4 min at 3,000 rpm, at 4°C) in the presence of a protease inhibitor cocktail (Roche). Aliquots were resuspended in SEM buffer (250 mM sucrose, 1 mM EDTA, and 10 mM MOPS-KOH, pH 7.2) and stored at −80°C.

### Submitochondrial assay

Membrane fractions were prepared by sonication of purified mitochondria. Purified mitochondria (50–500 μg) were resuspended in 1 mL SEM buffer containing 500 mM NaCl and sonicated on ice for 3 × 30 s. Samples were then centrifuged at 100,000 × g for 1 h at 4°C. The supernatant was collected as the soluble fraction. The pellet was solubilized in digitonin buffer (20 mM Tris-HCl, pH 7.4, 50 mM NaCl, 0.1 mM EDTA, 10% glycerol, and 1% digitonin (Calbiochem)) for 30 min on ice. The samples were clarified by centrifugation at 21,130 × g for 30 min at 4°C, after solubilization and concentrated. The soluble fraction was buffer-exchanged to remove excess salt. Protein concentrations were determined using a BCA protein assay (Thermo).

### SDS-PAGE, Blue and Clear Native PAGE and Western Blotting

Native protein complexes were separated by blue native PAGE (BN-PAGE) or clear native PAGE (CN-PAGE). Mitochondria were solubilized for 30 min on ice in solubilization buffer (20 mM Tris-HCl, pH 7.4, 50 mM NaCl, 0.1 mM EDTA, 10% glycerol, and 1% digitonin (Calbiochem)) followed by a centrifugation at 21,130 × g for 30 min at 4°C. For BN-PAGE, the supernatants were supplemented with BN loading dye (0.5% Coomassie blue G-250 and 10 mM Bis-Tris, pH 7.0) and loaded onto 3–12% polyacrylamide gradient gels (NativePAGE, Invitrogen). Electrophoresis was performed using BN anode buffer (50 mM Bis-Tris and 50 mM Tricine, pH 7.0) and BN cathode buffer (50 mM Bis-Tris and 50 mM Tricine, pH 7.0, 0.05% Coomassie blue G-250). For CN-PAGE, a similar protocol was used but omitting the Coomassie in the loading dye. For SDS-PAGE, samples were mixed with an equal volume of 2× Laemmli buffer containing 25 mM 2-mercaptoethanol and incubated for 10 min at 70°C. Denatured samples were loaded onto 4–12% polyacrylamide gels (NuPAGE, Invitrogen) and electrophoresis was run in MES/SDS running buffer (50 mM Bis-Tris, 50 mM Tris base, pH 7.3, 0.1% SDS, and 1 mM EDTA). For two-dimensional SDS-PAGE, strips cut from unstained BN-PAGE gels were loaded into the wells of 4–12% polyacrylamide 2D gels (Invitrogen) and run in MES/SDS running buffer. For western blotting, proteins were transferred from the gels to PVDF membranes using a dry transfer system (iBlot 2, Thermo). Membranes were stained with 1% Ponceau S and destained until protein bands were visible. The membranes were then cut as required and blocked overnight in blotting buffer (20 mM Tris-HCl, pH 8.0, 150 mM NaCl, and 0.05% Tween 20) supplemented with 3% BSA. Membranes were incubated with primary antibodies for 1 h at room temperature, washed, and incubated with a secondary anti-rabbit IgG antibody (Sigma; 1:15,000 dilution in blotting buffer). After washing, membranes were incubated in AP buffer (100 mM Tris-HCl, pH 9.5, 100 mM NaCl, and 5 mM MgCl2). CDP-Star (Thermo) was added, and the chemiluminescence signal was detected using a FusionCapt Advance FX7 16.11 system (Vilber Lourmat).

### Antibodies

Antibodies against all subunits but cyt *b* of yeast complex III were custom-generated against selected peptides. Anti-cyt *b* antibody (*21*) was a generous gift from Geneviève Dujardin and Brigitte Meunier (Université Paris-Saclay, CEA, CNRS, Institute for Integrative Biology of the Cell (I2BC), Gif-sur-Yvette, France).

### Mass Photometry

Mass photometry measurements were performed using a OneMP instrument (Refeyn Ltd.) with a controlled by AcquireMP software (v2.5; Refeyn Ltd.). Protein samples were diluted immediately before measurement. The flow chamber was filled with buffer, and the focus position was identified and maintained throughout each measurement using a feedback loop based on the total internal reflection of a reference laser beam. Approximately 15 µL of diluted sample was then introduced into the chamber, and data were recorded for 300 s. Protein-binding events at the glass surface were analyzed using DiscoverMP software (v2.5; Refeyn Ltd.). The instrument was calibrated using a mixture of protein standards with molecular masses of 66, 132, 335, and 669 kDa. Microfluidic mass photometry (MassFluidix) measurements were carried out using 1st-generation microfluidic chips (Refeyn, Ltd.). Heterologously expressed Qcr1/Qcr2 in 100mM Tris-HCl, 150mM NaCl, 1 mM EDTA pH 8.0, was pumped via the sample inlet. Inlet tubes were washed with water and primed with buffer before and between runs. Pressure and flow rates were monitored with the AcquireMP software. Histograms were created in DiscoverMP. MFP1 was used as mass calibrant.

### CryoEM grid preparation and data collection

Grids were vitrified using an EM GP2 automatic plunge freezer (Leica). Before sample application, holey-carbon cryo-EM grids were glow-discharged using an ELMO system (Cordouan Technologies; 2 mA, 0.2 mbar, 60 s). Grids were transferred to the EM GP2 chamber and approximately 3 µL of protein sample (3.5 mg/mL for recombinant Qcr1/Qcr2 complexes, 12 mg/ml for the sample purified from *crd1*Δ yeast cells) was applied to each grid. Grids were blotted for 3 s and plunge-frozen in liquid ethane. Initial electron-microscopy screening was performed at the BioEM Lab, University of Basel, Switzerland, using a Talos L120C microscope (Thermo Fisher Scientific), and subsequently at the Cryo-EM Facility Freiburg, Germany, using a Titan Krios G4 microscope equipped with a Falcon 4i direct electron detector (Thermo Fisher Scientific).

Datasets were collected at the Cryo-EM Facility Freiburg using the same microscope and detector as well as at the Centre for Integrative Biology (CBI), Institute of Genetics and of Molecular and Cellular Biology (IGBMC), Illkirch, France, using a Titan Krios equipped with a K3 direct electron detector (Gatan). Further datasets were collected at ESRF beamlines CM01 and CM02 (Grenoble, France), using a Titan Krios G3 equipped with a K3 detector and a Titan Krios G4 equipped with a Falcon 4i detector, respectively.

### CryoEM data processing

Beam-induced motion and stage drift were corrected by whole-frame alignment, and the contrast transfer function (CTF) was estimated using Unblur and CTFFIND, as implemented in cisTEM v1.0 or v2.0.0 beta (*22*) or using RELION 5.0 (*23*). Particles were automatically picked using crYOLO 1.7.5 (*24*). The crYOLO-picked particle positions were imported into cisTEM. and 2D classification was performed. and the resulting particle coordinates were imported into cisTEM for 2D classification. An initial 3D reconstruction was generated from the particles retained after 2D classification. Further refinement was performed using 3D auto-refine in cisTEM (*22*). The dataset was then imported into RELION 5.0 (*23, 25*) for CTF refinement and 3D refinement. Where necessary, 3D classification was used to reduce sample heterogeneity. Bayesian polishing and additional CTF refinement were then performed to generate the final maps. Final maps for all datasets were further improved by density modification using phenix.resolve_cryo_em (phenix1.19.2) (*26*). Density modification was performed using the two half-maps and a soft mask to exclude bulk-solvent noise. The resulting maps were used for model building and refinement.

### Model building and refinement of cryo-EM data

An initial structure model was obtained by manually fitting the individual chains from the X-ray crystallography structure of cytochrome *bc*_1_ (see below) into the obtained cryo-EM map using Coot (*27*). The atomic model was subsequently refined using phenix.real_space_refine (*26*). Hydrogen atoms were added to the coordinates using phenix.ready_set (*26*) and the resulting model was further refined with phenix.real_space_refine and manually rebuilt in Coot to improve the geometry. Model geometry was further improved using ISOLDE 1.11 (*28*), followed by a final refinement in phenix.real_space_refine. The final structures were validated using phenix.validation_cryoem (*26*). Figures were prepared using the Open Source PyMol 3.1 Molecular Graphics System (Schrödinger, LLC; NY, USA) and UCSF ChimeraX 1.11 (*29*).

### Crystallization of *bc*_1_ and X-ray diffraction data collection

Antibody-fragment mediated crystallization of wild type yeast complex III from strain BY4741 was performed essentially as described earlier (*20*) and with stigmatellin added as specific inhibitor. Large crystals could be obtained, cryo-protected and frozen in liquid nitrogen prior data collection. X-ray diffraction data were collected on beamline P13 operated by EMBL Hamburg at the PETRA III storage ring (DESY, Hamburg, Germany) (*30*). Data were processed using Autoproc (*31*) and data quality was assessed using Phenix.Xtriage (*26*). The phase problem was solved by molecular replacement using Phaser (*27*) and the yeast complex III structure (PDB ID 2IBZ) as search model. The model was rebuild in Coot (*32*) and refined using Buster (*33*) and Phenix (*26*). Figures were prepared using the Open Source PyMol 3.1 Molecular Graphics System (Schrödinger, LLC; NY, USA) and UCSF ChimeraX 1.11 (*29*).

### Mitochondrial network analysis in yeast

Yeast strains were grown at 30 °C in synthetic medium lacking tryptophan (0.17% yeast nitrogen base, 0.5% ammonium sulfate) with 2% glucose (SD-Trp) or 2% glycerol (SG-Trp). Overnight SD-Trp precultures were diluted into SD-Trp or, after two washes, into SG-Trp, kept in exponential phase and imaged after 24 h. Cells were stained in their growth medium with MitoTracker Green for 30 min at 30 °C, washed once and immobilised on concanavalin A-coated glass-bottom dishes. Z-stacks of whole cells (0.25 µm steps) were acquired at room temperature on a DeltaVision OMX Flex microscope (UPlanSApo ×60/1.4 oil Olympus objective, PCO Edge 4.2 sCMOS camera) with identical settings for all conditions, from fields selected in brightfield, and deconvolved in softWoRx. Mitochondrial networks of single cells were quantified in 3D with MitoGraph (*34*). Data are shown as box plots (median, interquartile range, whiskers 1.5× IQR) and were compared by one-way ANOVA with Sidak’s multiple comparisons test in R (emmeans).

### Complexome profiling

Complexome profiling was performed essentially as described (*35*).

## Acknowledgments

We acknowledge Carola Alampi and Mohamed Chami (BioEM lab, University of Basel. Switzerland) for assistance during initial screening of cryo-EM samples. We acknowledge the expertise and resources of the French Infrastructure for Integrated Structural Biology (FRISBI) ANR-10-INBS-0005 and thank Corinne Crucifix, Alexandre Durand and Nils Marechal (CBI, IGBMC, Illkirch, France) for assistance during cryo-EM data acquisition. Similarly, electron microscopy data were collected at the Cryo-EM Facility of the University of Freiburg (RRID: SCR_025860). The Titan Krios G4 cryo-TEM used for imaging was funded by Deutsche Forschungsgemeinschaft (project no. 506518771) and is operated within the Microscopy and Image Analysis Platform (MIAP), University of Freiburg. We acknowledge the European Synchrotron Radiation Facility (ESRF) for provision of beamtime under proposal ID MX-2365 and MX-2702 and on beamlines CM01. We thank the staff of the ESRF and EMBL Grenoble for assistance and support in using the jointly operated Structural Biology beamlines. Finally, we thank the staff of beamline P13 operated by EMBL Hamburg at the PETRA III storage ring (DESY, Hamburg, Germany) for the support during X-ray diffraction data collection. We are grateful to Geneviève Dujardin and Brigitte Meunier (Université Paris-Saclay, CEA, CNRS, Institute for Integrative Biology of the Cell (I2BC), Gif-sur-Yvette, France) for the generous gift of an antibody for cyt *b*.

Artificial intelligence (AI)-based language tools (Claude, Anthropic, and ChatGPT, OpenAI) were used to assist with wording and language refinement. These tools were not used to generate scientific content, interpret results or draw conclusions. After using these tools, the authors reviewed, verified and edited the content as needed and take full responsibility for the content of the publication.

## Funding

Deutsche Forschungsgemeinschaft (DFG, German Research Foundation) under Germany’s Excellence Strategy, CIBSS – EXC-2189 – Project ID 390939984 (CH, CK, PH, NW)

Deutsche Forschungsgemeinschaft Project-ID 403222702 – SFB 1381 (CH, CK, PH, NW)

Deutsche Forschungsgemeinschaft Project-ID 431984000 – SFB 1453 (CH)

Deutsche Forschungsgemeinschaft Project-ID 423813989 – GRK 2606 (CK)

Deutsche Forschungsgemeinschaft Project-IDs 450216812 and 409673687 (CK)

European Union’s Horizon Europe research and innovation programme, Grant Agreement No 101224197 (CK). Funded by the European Union. Views and opinions expressed are however those of the author(s) only and do not necessarily reflect those of the European Union or the European Research Council Executive Agency (ERCEA). Neither the European Union nor the granting authority can be held responsible for them.

## Author contributions

**Jorge Jimenez-Niebla:** Conceptualization, Methodology, Investigation, Visualization, Writing – original draft, Writing – review & editing; **Roshan Jha:** Methodology, Investigation, Visualization; **Christophe Wirth:** Conceptualization, Methodology, Investigation, Visualization, Writing – original draft, Writing – review & editing; **Jon V. Busto:** Investigation, Visualization; **Antoine Feignier:** Investigation, Visualization; **Miguel Cosenza-Contreras:** Investigation, Visualization; **Katharina Ruoff:** Investigation, Visualization; **Manuel Dorer:** Investigation, Visualization; **Riccardo Babic:** Investigation, Visualization; **Friedel Drepper:** Methodology, Investigation, Visualization; **Claudine Kraft:** Conceptualization, Supervision, Funding acquisition, Writing – original draft, Writing – review & editing; **Pitter F. Huesgen:** Conceptualization, Supervision, Funding acquisition, Writing – original draft, Writing – review & editing; **Nils Wiedemann:** Conceptualization, Supervision, Funding acquisition, Writing – original draft, Writing – review & editing; **Wei-Chun Kao:** Conceptualization, Methodology, Investigation, Visualization, Supervision, Writing – original draft, Writing – review & editing; **Carola Hunte:** Conceptualization, Methodology, Investigation, Visualization, Supervision, Writing – original draft, Writing – review & editing;

## Data, code, and materials availability

All data are available in the main text or the supplementary materials. Atomic coordinates of the reported cryo-EM and X-ray structures and the cryo-EM maps are in preparation for deposition in the Protein Data Bank (PDB) and the Electron Microscopy Data Bank (EMDB) and will be made publicly available upon peer-reviewed publication. Accession codes will be provided prior to formal publication.

