## Supplemental Data for "Modular assembly of mitochondrial complex III from a dynamic matrix domain pool"

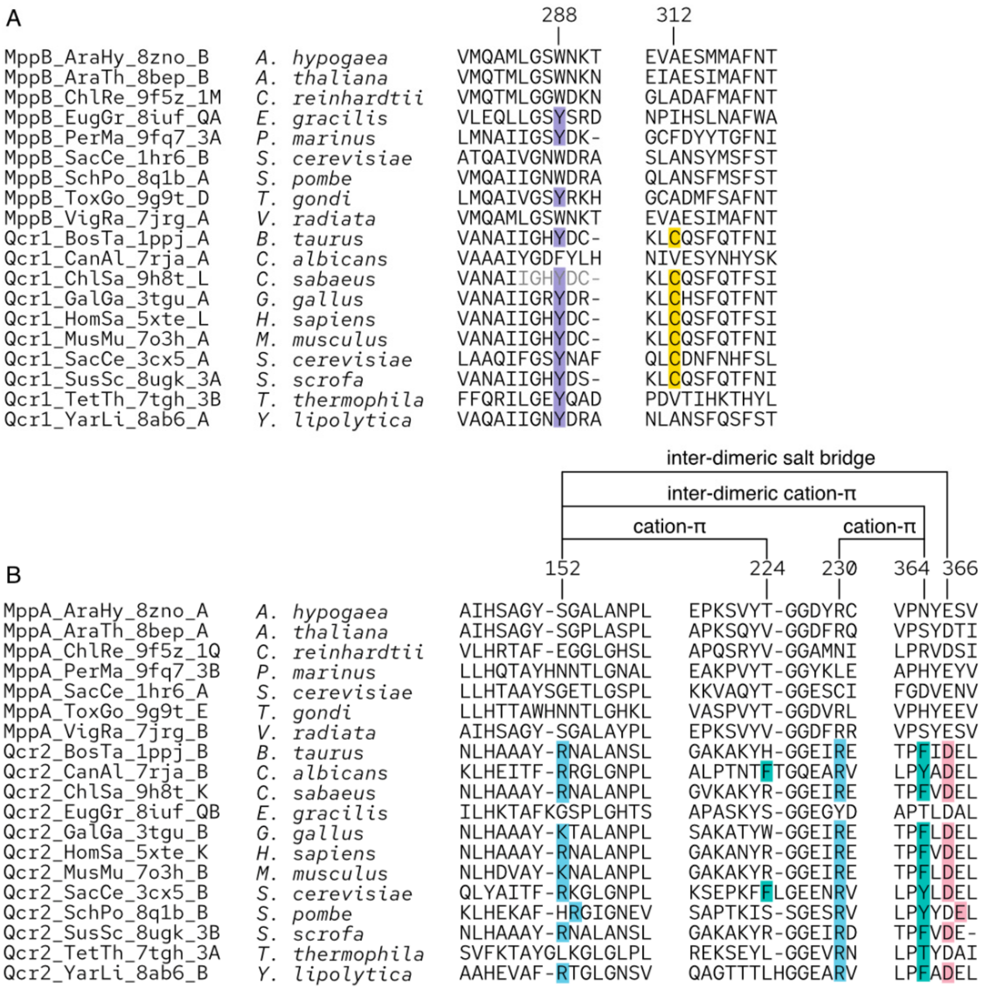


**Fig. S1:** Multiple structural alignment of interface residues in Qcr1 (A) and Qcr2 (B). Structural alignment was performed using FoldMason (*36*) with X-ray and cryo-EM structures of respiratory chain complexes which contain Qcr1, Qcr2, MppB or MppA as subunits. The sequences are annotated using the pattern of protein name_species abbreviations_pdb code_chain ID. For Qcr2, inter-dimeric interactions are established through one amino acid residue from one protomer and the other residue from the two-fold symmetry-related protomer.

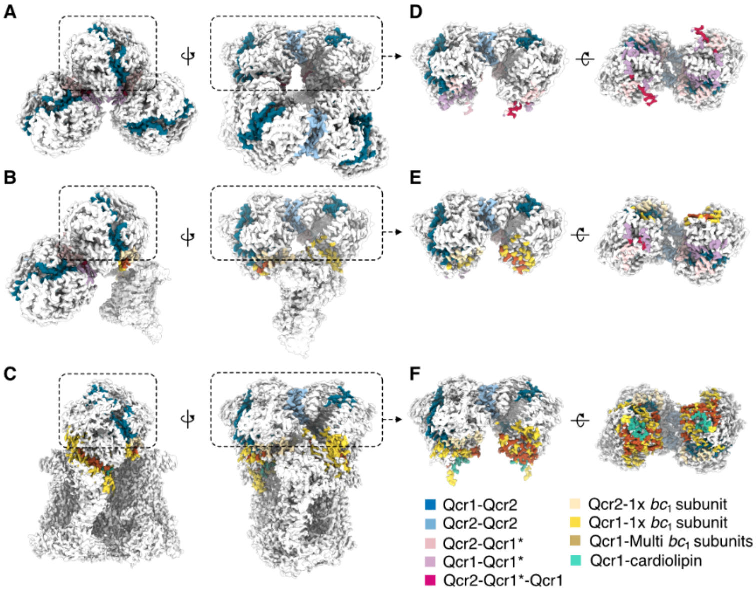


**Fig. S2:** Interfaces of Qcr1 and Qcr2 in $CM_{Hexa}$ *crd1*Δ (A), pre-dimerization intermediate (PDI) *crd1*Δ (B), and complex III (C, Ybc1) (for structures see Table S1). Interfaces were analysed using the EMBL-EBI PISA server (37) and visualised in ChimeraX. The subunits in the dashed-rectangular box are shown in panels D, E, and F in order to reveal the interfaces. Qcr1* denotes the Qcr1 not shown in panels D and E.

**Table S1:** Cryo-EM and X-ray structures determined for this study. cryo-EM: cryogenic electron microscopy; XRD: X-ray crystallography. Resolution is specified in Å.

| Structure | Method | Å | Comment |
|---|---|---|---|
| **$CM_{Hexa}$ *crd1Δ*** | cryo-EM | 2.46 | Core module hexamer obtained from CRD1Δ |
| **PDI *crd1Δ*** | cryo-EM | 2.67 | Pre-Dimerization Intermediate obtained from CRD1Δ |
| **$CM_{Hexa}$ RC** | cryo-EM | 2.80 | Recombinant "wild-type" core module hexamer |
| **$CM_{Tetra}$ RC** | cryo-EM | 3.20 | Recombinant "wild-type" core module tetramer |
| **$CM_{Duo}$ RC** | cryo-EM | 3.61 | Recombinant "wild-type" core module dimer |
| **$CM_{Octa}$ ΔCT** | cryo-EM | 3.80 | Recombinant "Qcr1 C-terminal truncated" core module octamer |
| **$CM_{Hexa}$ ΔCT** | cryo-EM | 3.26 | Recombinant "Qcr1 C-terminal truncated" core module hexamer |
| **$CM_{Tetra}$ ΔCT** | cryo-EM | 3.62 | Recombinant "Qcr1 C-terminal truncated" core module tetramer |
| **$CM_{Mono}$ R152A** | cryo-EM | 3.33 | Recombinant "Qcr2 R152A" core module monomer |
| **$CM_{Mono}$ R152W** | cryo-EM | 3.54 | Recombinant "Qcr2 R152W" core module monomer |
| **$CM_{Hexa}$ Y288A** | cryo-EM | 2.16 | Recombinant "Qcr1 Y288A" core module hexamer |
| **$CM_{Tetra}$ Y288A** | cryo-EM | 2.28 | Recombinant "Qcr1 Y288A" core module tetramer |
| **$CM_{Duo}$ Y288A** | cryo-EM | 2.61 | Recombinant "Qcr1 Y288A" core module dimer |
| **$CM_{Hexa}$ Y288S** | cryo-EM | 2.58 | Recombinant "Qcr1 Y288S" core module hexamer |
| **$CM_{Tetra}$ Y288S** | cryo-EM | 2.78 | Recombinant "Qcr1 Y288S" core module tetramer |
| **$CM_{Duo}$ Y288S** | cryo-EM | 3.40 | Recombinant "Qcr1 Y288S" core module dimer |
| **Ybc1** | XRD | 1.90 | Yeast cyt $bc_1$ complex co-crystalized with Fv antibody fragment |
